# TGF-β-dependent Lymphoid Tissue Residency of Stem-like T cells Limits the Response to Tumor Vaccine

**DOI:** 10.1101/2021.07.19.452945

**Authors:** Guo Li, Liwen Wang, Chaoyu Ma, Wei Liao, Yong Liu, Shruti Mishra, Xin Zhang, Yuanzheng Qiu, Qianjin Lu, Nu Zhang

**Author notes:** Corresponding author: N.Z.

## Abstract

Stem-like CD8^+^ T cells represent the key subset responding to multiple tumor immunotherapies, including tumor vaccination. However, the signals that control the differentiation of stem-like T cells are not entirely known. Most previous investigations on stem-like T cells are focused on tumor infiltrating T cells (TIL). The behavior of stem-like T cells in other tissues remains to be elucidated. Tissue-resident memory T cells (T_RM_) are often defined as a non-circulating T cell population residing in non-lymphoid tissues. TILs carrying T_RM_ features are associated with better tumor control. Here, we found that stem-like CD8^+^ T cells differentiated into T_RM_s in a TGF-β and tumor antigen dependent manner almost exclusively in tumor draining lymph node (TDLN). TDLN-resident stemlike T cells were negatively associated with the response to tumor vaccine. In other words, after tumor vaccine, TDLN stem-like T cells transiently lost T_RM_ features, differentiated into migratory effectors and exerted tumor control.

## Introduction

Tumor immunotherapy represents one of the most prominent biomedical advances in recent decades(*1–3*). Some tumor immunotherapies, such as immune checkpoint blockade and tumor vaccines aim at reinvigorating endogenous T cells, including tumor-antigen-specific CD8^+^ T cells. Persistent antigen exposure (e.g., tumor antigen) induces T cell exhaustion with reduced effector function(*4*). Exhausted CD8^+^ T cells are heterogenous and a less exhausted subset carries stem cell-like features(*5–9*). These stem-like CD8^+^ T cells express transcription factor Tcf-1 (T cell factor-1) and sustain CD8^+^ response during chronic antigen exposure. Importantly, these stem-like CD8^+^ T cells are the ones responding to immune checkpoint blockade therapies(*6, 8, 10*) and correlating with the efficacy of tumor vaccines(*11, 12*). However, the signals that control the maintenance, differentiation and migration of these stem-like T cells are not entirely known.

Transforming growth factor-β (TGF-β) is a pleiotropic cytokine. When focusing on antitumor immunity, TGF-β is generally considered as an immune suppressor. It is well known that TGF-β directly inhibits the proliferation and effector function of CD8^+^ T cells(*13–19*). Systemic blocking TGF-β synergizes with tumor vaccine or PD-1/PD-L1 blockade to boost CD8^+^ T cell response in mouse models (*20–24*). Recent publications have established that blocking TGF-β signaling can boost the efficacy of PD-1/PD-L1 blockade therapy by targeting tumor stromal compartment(*25–28*). In addition, suppression of CD4^+^ T cell-intrinsic TGF-β signaling promotes tumor control via regulating type 2 immunity mediated blood vasculature remodeling(*29, 30*). Because most studies about TGF-β on tumorspecific CD8^+^ T cells were carried out without incorporating the knowledge of stem-like T cells, it is critical to revisit the function of TGF-β on tumor-specific CD8^+^ T cells, especially on stem-like CD8^+^ T cells.

Tissue-resident memory T cells (T_RM_) represent a unique memory T cell population, which is separated from the circulation and maintained in a self-sustained manner(*31–34*). Originally discovered in acute infection models, T_RM_ has been established as an essential component of tissue-specific immunity. Surprisingly, recent findings have demonstrated that stem-like CD8^+^ T cells generated after chronic viral infection bear similar properties as T_RM_, i.e., mostly confined to secondary lymphoid organs (e.g., spleen and lymph nodes) and non-circulating(*35, 36*). Thus, residency inside lymphoid tissues may be essential for stem-like T cells. However, whether similar scenario exists in tumor immunity settings remains unknown. The signals that control the T_RM_ properties of stem-like T cells remain to be defined. Whether there is any functional connection between T_RM_-like and stem-like features for this unique T cell subset remains unexplored. The vast majority of previous research on tumor-specific CD8^+^ T cells, including stem-like T cells is focused on TILs. Even though the cancer-immunity cycle model(*37*) is widely accepted, tumor-specific CD8^+^ T cells residing in lymphoid organs is substantially underappreciated.

It is well known that TGF-β signaling to CD8^+^ T cells is essential for the differentiation and maintenance of T_RM_ after acute infection(*33–43*). When focusing on tumor immunity, T_RM_-like signature has often been positively associated with the capacity of TILs to control tumor(*44–51*). It remains a mystery how to reconcile the facts that TGF-β promotes T_RM_, T_RM_ limits tumor growth and TGF-β blockade improves tumor control.

Here, to revisit the role of TGF-β in tumor-specific CD8^+^ T cells, especially stem-like T cells, we employed B16F10 transplanted melanoma model together with B16 tumorspecific Pmel-1 TCR transgenic mice as well as B16-OVA and OT-1 system. We discovered that TGF-β receptor deficient tumor-specific CD8^+^ T cells exhibited dramatically enhanced anti-tumor activity only after tumor vaccination. Importantly, the superior response to tumor vaccine in TGF-βR deficient CD8^+^ T cells was completely abolished when the migration of T cells from tumor draining lymph nodes (TDLN) to tumor was inhibited. When TDLN was examined, we found that a significant portion of stem-like CD8^+^ T cells differentiated into T_RM_ in a TGF-β-dependent manner. Tumor vaccine induced dramatic loss of T_RM_ features in stem-like T cells isolated from TDLN. Together, TDLN functions as a resident reservoir of stem-like T cells. Loss of T_RM_ phenotype is required for the differentiation of migratory effectors and superior response to tumor vaccine.

## Results

### Disruption of TGF-β receptor in tumor-specific CD8^+^ T cells alone is not sufficient for tumor control

To examine the role of TGF-β signaling in stem-like T cells during tumor immunotherapies, we generated mature T cell-specific TGF-β receptor conditional knockout mice (*Tgfbr2*^f/f^ distal *Lck*-Cre, hereafter referred to as *Tgfbr2^-/-^*)(*13*). Further, we have established WT (wild type) and *Tgfbr2^-/-^* Pmel-1 TCR transgenic mice carrying CD8^+^ T cells specific for an endogenous melanocyte epitope (gp100_25-33_ presented by H-2D^b^ (*52*)). To be noted, gp100_25-33_ represents an endogenous tumor antigenic peptide derived from B16 melanoma. All of our Pmel-1 mice (both WT and *Tgfbr2^-/-^*) carried congenic markers so that donor Pmel-1 T cells could be easily followed after adoptive transfer.

To mimic endogenous anti-tumor immunity, we adoptively transferred naïve Pmel-1 T cells (10^5^ cells/mouse) into each unmanipulated WT mouse before tumor inoculation via a subcutaneous (s.c.) route. To determine the efficacy of anti-tumor responses, naïve WT and *Tgfbr2^-/-^* Pmel-1 T cells were separately transferred into different recipients (**Fig. 1A**). Surprisingly, we did not observe significant difference in tumor control between recipients of naïve WT vs *Tgfbr2^-/-^* Pmel-1 T cells (**Fig. 1B** and **1C**). Further, we repeated the experiments with OT-1 TCR transgenic mice recognizing a peptide derived from chicken ovalbumin (OVA) and a B16-OVA tumor line. WT OT-1 and *Tgfbr2^-/-^* OT-1 exhibited similar anti-tumor immunity in WT hosts (**Fig. S1A** and **S1B**). These findings are consistent with the recent publications, which emphasize on the impacts of TGF-β signaling on CD4^+^ T cells(*29, 30*) and tumor stroma(*25–28*). Our result suggests that suppression of TGF-β signaling in naturally primed CD8^+^ T cells alone is not sufficient to control aggressive tumors, such as B16 melanoma.

**Figure 1.**
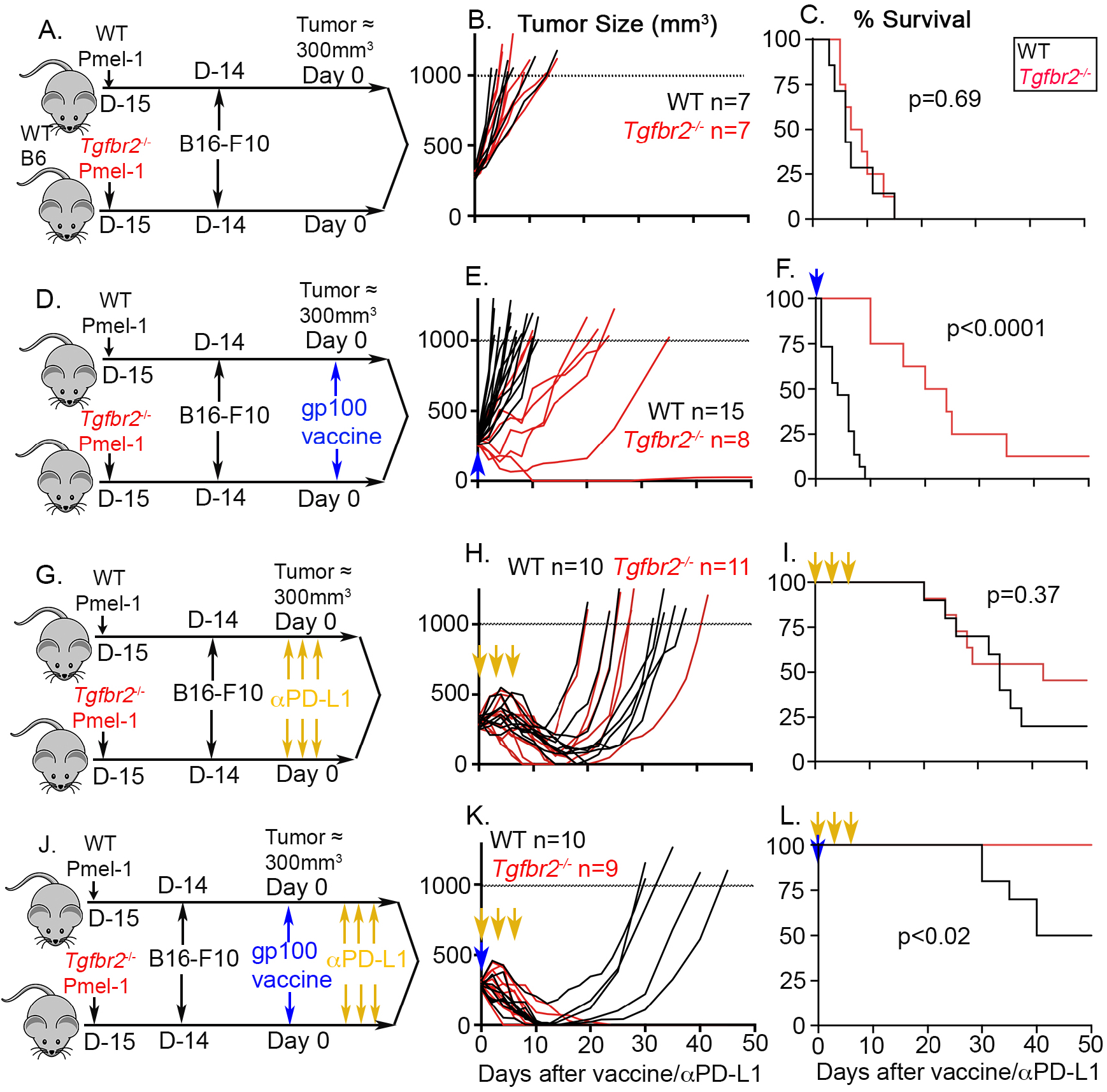
*Tgfbr2^-/-^* CD8^+^ T cells exhibited greatly enhanced responses to tumor vaccine. (**A**) to (**C**), Pmel-1 alone. (**A**) Schematics; (**B**) tumor growth; (**C**) Survival curve. (**D**) to (**F**), Pmel-1 +tumor vaccine. (**D**) Schematics; (**E**) tumor growth; (**F**) Survival curve. (**G**) to (**I**), Pmel-1 +αPD-L1. (**G**) Schematics; (**H**) tumor growth; (**I**) Survival curve. (**J**) to (**L**), Pmel-1+tumor vaccine+ αPD-L1. (**J**) Schematics; (**K**) tumor growth; (**L**) Survival curve. Pooled results from 2-3 independent experiments for each setting. Each line in (B), (E), (H) and (K) represents the results from an individual mouse. *P* values were calculated by Mantel-Cox test.

### *Tgfbr2^-/-^* CD8^+^ T cells synergize with tumor vaccine but not with PD-L1 blockade

Next, we s.c. administrated a tumor vaccine [Poly I:C+gp100_25-33_ (Pmel-1 cognate peptide)] once when tumor size reached around 300mm^3^ (**Fig. 1D**). Most likely due to the facts that tumor vaccine was given at a relatively late stage (around 300mm^3^), vaccine did not boost anti-tumor immunity in the recipients of WT Pmel-1 T cells (compare black lines in **Fig. 1B** vs **1E** and **Fig. 1C** vs **1F**). In contrast, *Tgfbr2^-/-^* Pmel-1 cells elicited significantly improved tumor control after vaccination **(Fig. 1E** and **1F**). Similar results were observed when B16-OVA and OT-1 system was used, i.e., *Tgfbr2^-/-^* OT-1 exhibited significantly improved tumor control after administration of OVA peptide vaccine (**Fig. S1C** to **S1E**).

In contrast, PD-L1 blocking antibody boosted the anti-tumor immunity of both WT and *Tgfbr2^-/-^* Pmel-1 T cells to a similar extent (compare **Fig. 1B** vs **1H** and **1C** vs **1I**). In other words, there was no significant difference between WT and *Tgfbr2^-/-^* Pmel-1 T cells after PD-L1 blockade (**Fig. 1H** and **1I**). Finally, we combined αPD-L1 blocking antibody with tumor vaccine (**Fig. 1J**). Under this setting, WT Pmel-1+ vaccine+αPD-L1 exhibited significantly improved response with 50% of mice cured of tumor. In contrast, 100% of *Tgfbr2^-/-^* Pmel-1+vaccine+αPD-L1 treated animals rapidly irradicated late-stage tumors (**Fig. 1K** and **1L**).

Together, we have demonstrated that *Tgfbr2^-/-^* CD8^+^ T cells exhibit superior response to tumor vaccine. Vaccine activated CD8^+^ T cells further synergize with PD-L1 blockade therapy. Because the most striking difference between WT and *Tgfbr2^-/-^* T cells was induced only after tumor vaccine, we would primarily focus on the response to tumor vaccine in the following studies.

### Greatly enhanced accumulation of *Tgfbr2^-/-^* effector T cells in tumor after vaccination

To determine the mechanisms underlying improved response to tumor vaccine for *Tgfbr2^-/-^* T cells, we first focused on tumor infiltrating Pmel-1 T cells before and after tumor vaccine administration (illustrated in **Fig. 2A**). Indeed, tumor vaccine significantly boosted the total number of *Tgfbr2^-/-^* Pmel-1 T cells in TILs (**Fig. 2B**). In contrast, even though there was a similar trend, the difference of WT Pmel-1 T cells before vs after vaccination did not reach statistical significance (**Fig. 2B**). Interestingly, the stem-like subset of TIL Pmel-1 T cells was significantly reduced for *Tgfbr2^-/-^* cells before vaccination. Both WT and *Tgfbr2^-/-^* Pmel-1 T cells carried a similarly reduced stem-like subset after vaccination (**Fig. 2C**). When B16-OVA and OT-1 system was examined, *Tgfbr2^-/-^* OT-1 T cells exhibited significantly enhanced accumulation inside tumor after vaccination (**Supp. Fig. 2B**). No significant difference of Tcf-1^+^ stem-like T cells were detected between WT vs *Tgfbr2^-/-^* OT-1 T cells before or after vaccination (**Supp. Fig. 2C**).

**Figure 2.**
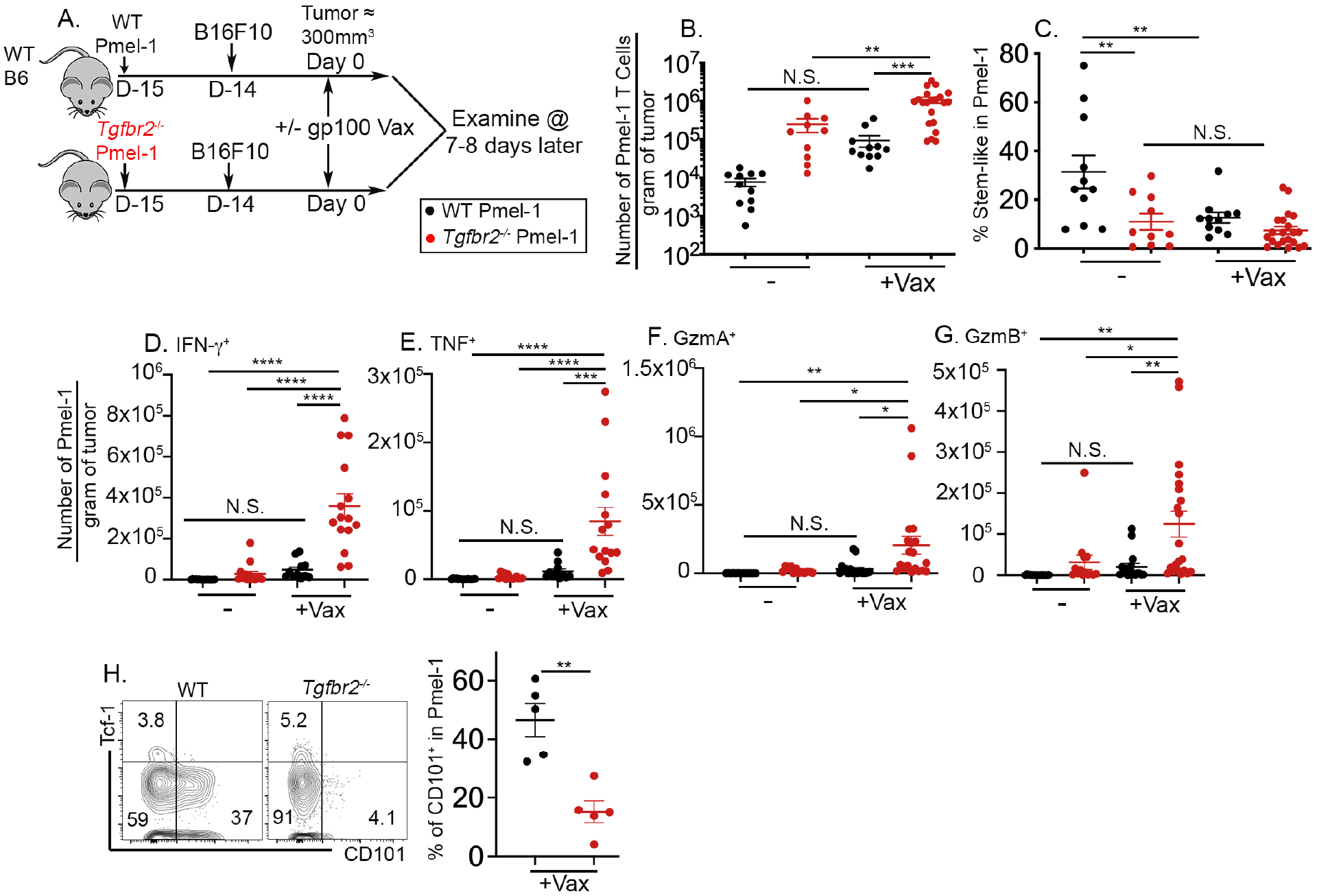
Dramatically increased accumulation of *Tgfbr2^-/-^* effector CD8^+^ T cells inside tumor after vaccination. (**A**) Experimental design. (**B**) The number of donor Pmel-1 T cells per gram of tumor is shown. (**C**) The percentage of stem-like subset in Pmel-1 T cells isolated from tumor is shown. Per gram of tumor, the numbers of donor Pmel-1 T cells producing IFN-γ (**D**), TNF (**E**), Granzyme A (**F**) and Granzyme B (**G**) are shown. (**H**) Left, representative FACS profiles of pre-gated donor Pmel-1 T cells isolated from tumor are shown; Right, the percentage of CD101^+^ subset in Pmel-1 T cells is shown. Each symbol represents the results from an individual mouse. Pooled results from 2 (H) or 3 (B to G) independent experiments are shown. N.S., not significant, *, p<0.05, **, p<0.01, ***, p<0.001 and ****, p<0.0001 by Student *t*-test (H) or Ordinary one-way ANOVA with multi-comparison posttest (B to G).

We further confirmed that only after tumor vaccine, TIL *Tgfbr2^-/-^* Pmel-1 T cells exhibited greatly enhanced effector functions, including the production of IFN-γ, TNF, granzyme A and granzyme B (**Fig. 2D** to **2G**). Consistently, tumor infiltrating *Tgfbr2^-/-^* Pmel-1 T cells (**Fig. 2H**) and *Tgfbr2^-/-^* OT-1 T cells (**Supp. Fig. 2D**) carried dramatically reduced population of terminally exhausted CD101^+^ subset after vaccination(*53, 54*). Together, enhanced tumor control (**Fig. 1**) is likely due to greatly enhanced accumulation of *Tgfbr2^-/-^* effector T cells infiltrating tumor after vaccination.

### Cell migration is required for *Tgfbr2^-/-^* Pmel-1 T cells to respond to tumor vaccine

It has been shown that stem-like T cells are the ones responding to tumor vaccine(*11, 12*). We had been puzzled by our findings that *Tgfbr2^-/-^* TILs carried less (**Fig. 2C**) or similar (**Supp. Fig. 2C**) stem-like subset before tumor vaccine and exhibited enhanced response to tumor vaccine (**Fig. 1**,**2** and **Supp. Fig. 1**,**2**). Considering widely accepted cancer-immunity cycle(*37*), we wondered whether active communication between lymphoid organs and tumors was involved in the response to tumor vaccine under our settings. To this end, we employed FTY720, a small molecule targeting S1PR1 (Sphingosine-1-phosphate Receptor 1) and inhibiting T cell egress from lymphoid organs. We started FTY720 treatment one day before vaccination to ensure stable concentration of FTY720 reached inside experimental animals (illustrated in **Fig. 3A**). Consistently, tumor vaccine elicited greatly enhanced tumor control in mice received *Tgfbr2^-/-^* Pmel-1 T cells. FTY720 treatment completely abolished this response (**Fig. 3B**). Further, FTY720 significantly enhanced the population of *Tgfbr2^-/-^* Pmel-1 T cells inside tumor draining lymph node (TDLN) (**Fig. 3C**) while abolished the increase of *Tgfbr2^-/-^* tumor infiltrating Pmel-1 T cells after vaccination (**Fig. 3D**). This result clearly demonstrates that cell migration from TDLN to tumor is essential for *Tgfbr2^-/-^* Pmel-1 T cells to respond to tumor vaccine.

**Figure 3.**
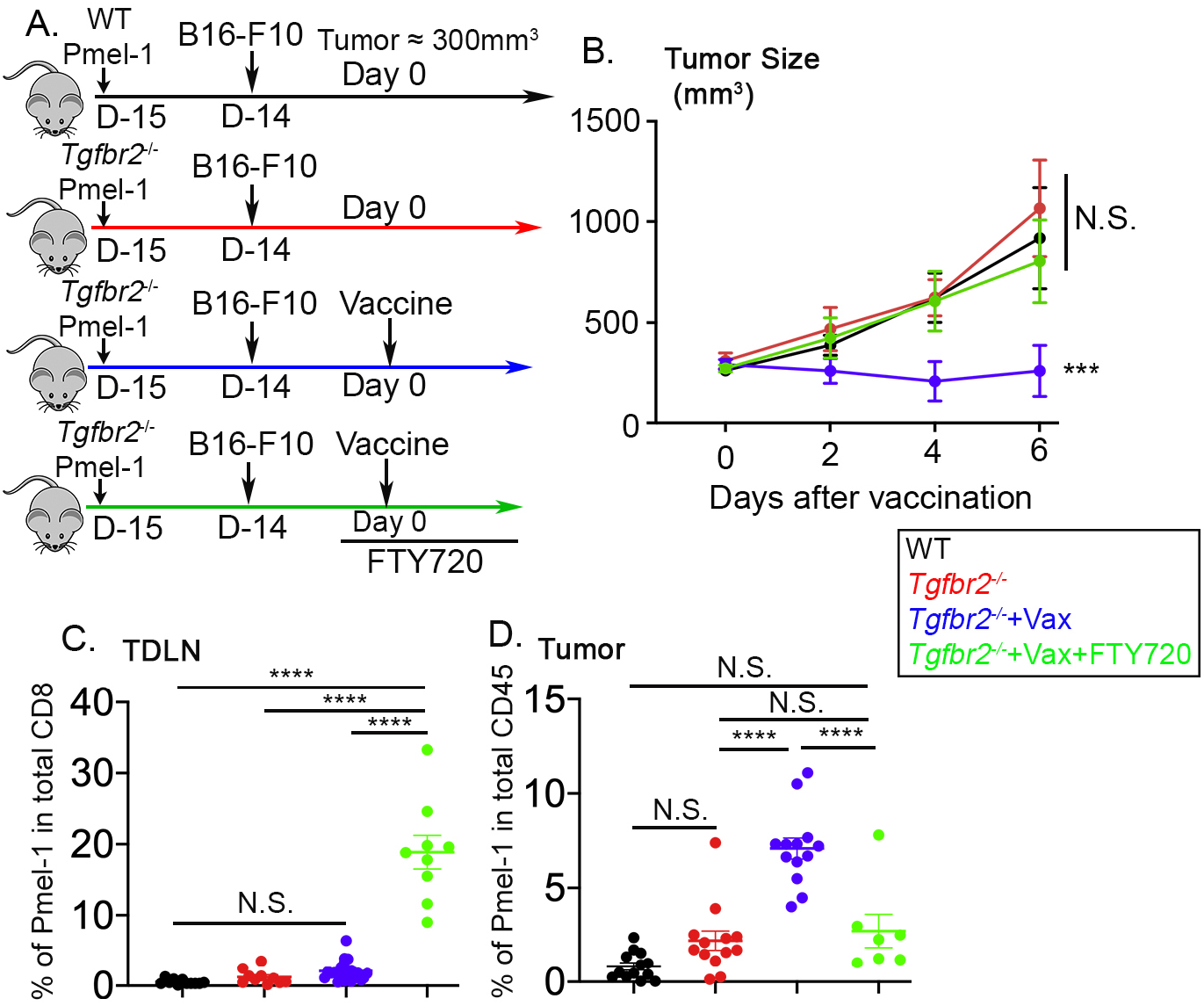
Cell migration is required for *Tgfbr2^-/-^* Pmel-1 T cells to respond to tumor vaccine. (**A**) Experimental design. (**B**) Tumor size (n=5-8 for each group). Statistical analysis was performed on day 6 results. (**C**) The percentage of donor Pmel-1 T cells in total CD8^+^ T cells isolated from TDLN is shown. (**D**) The percentage of donor Pmel-1 T cells in total CD45^+^ cells isolated from tumor is shown. Each symbol in (C) and (D) represents the results from an individual mouse. Pooled results from 3 independent experiments are shown. N.S., not significant, ***, p<0.001 and ****, p<0.0001 by Ordinary one-way ANOVA with multi-comparison posttest.

Further, it has been demonstrated that CX3CR1^+^ effector CD8^+^ T cells representing a migratory subset with enhanced effector functions(*55*). Indeed, we found that CX3CR1^+^ Pmel-1 T cells were undetectable before tumor vaccine in TDLN. Tumor vaccine greatly boosted CX3CR1^+^ subset in TDLN, especially in *Tgfbr2^-/-^* cells (**Supp. Fig. 3A** and **3B**), leading to dramatically increased accumulation of this effector subset in tumors (**Supp. Fig. 3C**). Together, tumor vaccine induces the differentiation of migratory effectors in TDLN and TGF-β suppresses this process.

### TGF-β and tumor antigen together induce T_RM_ phenotype on stem-like T cells in TDLN

It is well established that TGF-β provides an essential signal for T_RM_ differentiation after acute infections(*38–43*). Stem-like T cells largely reside inside secondary lymphoid organs without circulation during chronic LCMV (lymphocytic choriomeningitis virus) infection(*35*). Based on these facts and our findings that *Tgfbr2^-/-^* Pmel-1 T cells exhibited enhanced migration from TDLN to tumor after vaccination, we hypothesized that stemlike Pmel-1 T cells differentiate into T_RM_ inside TDLN in a TGF-β-dependent manner.

To test this hypothesis, we employed widely accepted T_RM_ marker CD69 and CD103. As shown in **Fig. 4A**, WT Pmel-1 T cells were adoptively transferred into B6 hosts followed by B16F10 tumor inoculation. After tumor grew to more than 400mm^3^, the distribution and phenotype of donor Pmel-1 T cells were examined. As expected, tumor-specific Pmel-1 T cells were highly enriched in TDLN (**Fig. 4B**). Interestingly, stem-like T cells were also enriched in TDLN (**Fig. 4C**). Other secondary lymphoid organs [including both nondraining LN (NDLN) and spleen] harbored less stem-like T cells than TDLN, but more than tumors (**Fig. 4C**). Importantly, CD69^+^CD103^+^ T_RM_ phenotype was largely restricted to stem-like T cells isolated from TDLN (**Fig. 4D** and **4E**).

**Figure 4.**
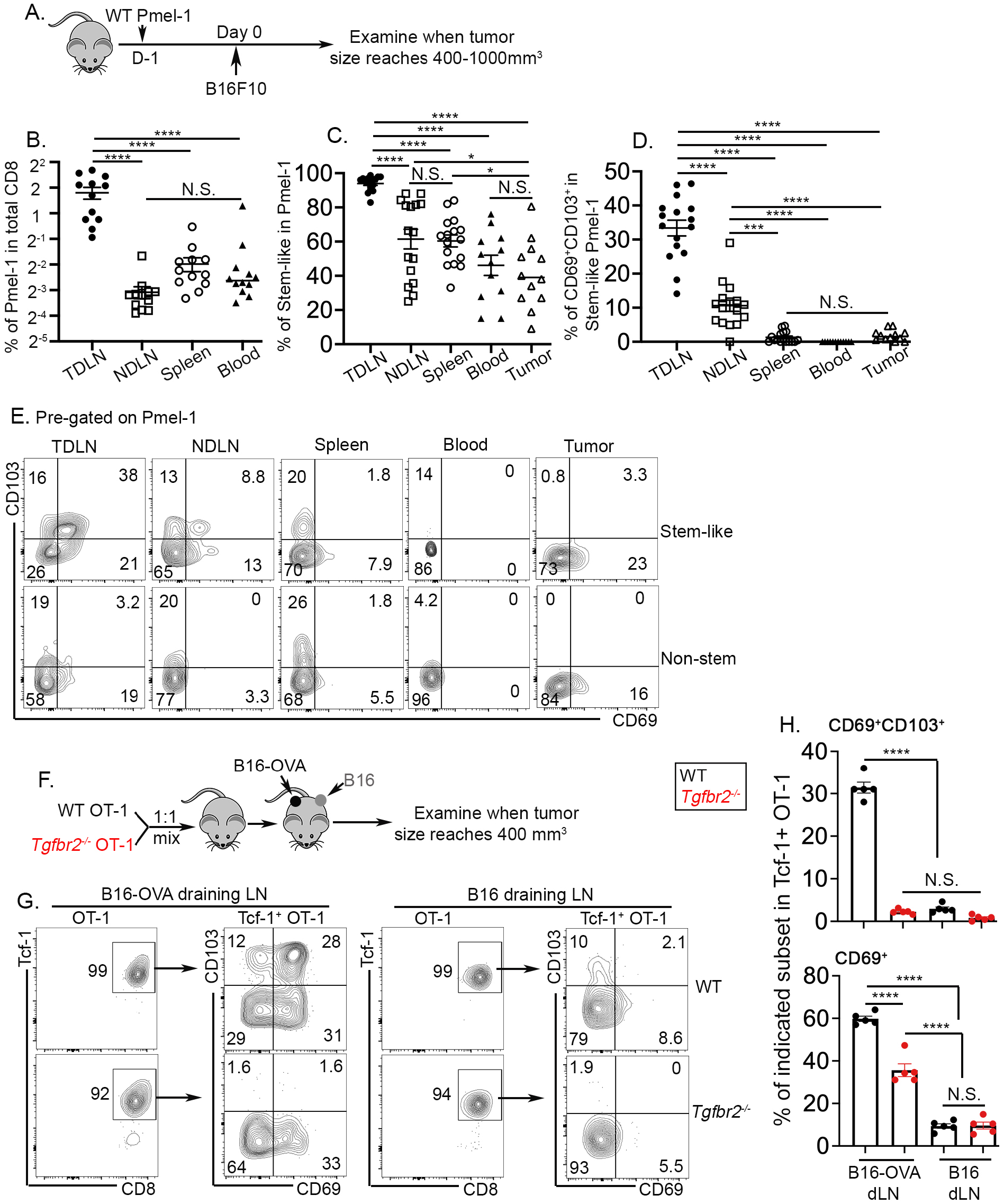
Tumor antigen and TGF-β-dependent establishment of T_RM_ stem-like T cells in TDLN. (**A**) Experimental design for (B) to (E). The percentage of Pmel-1 T cells in total CD8^+^ cells (**B**), the percentage of stem-like subset in Pmel-1 T cells (**C**) and the percentage of CD69^+^CD103^+^ cells in stem-like Pmel-1 T cells (**D**) isolated from different tissues of tumor bearing mice are shown. (**E**) Representative FACS profiles of stem-like (Upper) and non-stem (Lower) Pmel-1 T cells isolated from different tissues are shown. Pooled results from 4 independent experiments are shown in (A) to (E). (**F**) Experimental design for (G) and (H). (**G**) Representative FACS profiles of donor OT-1 T cells isolated from B16-OVA draining LN (Left) and B16 draining LN (Right) are shown. (**H**) The percentage of CD69^+^CD103^+^ cells in stem-like OT-1 T cells (Upper) and the percentage of CD69^+^ cells in stem-like OT-1 T cells (Lower) are shown. Each symbol represents the results from an individual recipient. N.S., not significant, *, p<0.05, ***, p<0.001 and ****, p<0.0001 by Ordinary one-way ANOVA with multi-comparison posttest.

To address the question why T_RM_-like stem-like T cells were enriched in TDLN, we examined the impacts of tumor antigen. To this end, we employed two B16 tumor lines, one with constitutive expression of model antigen OVA (B16-OVA) and one without (B16). As illustrated in **Fig. 4F**, WT and *Tgfbr2^-/-^* OT-1 were adoptively co-transferred into B6 hosts. All host mice were s.c. inoculated with B16 and B16-OVA on the opposite sides. In this system, the contribution from TGF-β and tumor antigen could be precisely determined in the same animal. As shown in **Fig. 4G** and **4H**, WT Tcf-1^+^ OT-1 differentiated into CD69^+^CD103^+^ cells in B16-OVA-draining LN, but not in B16-draining LN. The expression of CD103 was TGF-β-dependent. Consistent with the findings in acute infection-induced T_RM_ (*43, 56, 57*), the induction of CD69 was reduced, but not completely abolished in *Tgfbr2^-/-^* cells. Importantly, the highly restricted distribution of CD69^+^CD103^+^ OT-1 in B16-OVA-draining LN, but not in B16-draining LN demonstrated that CD69^+^CD103^+^ cells were bona fide T_RM_ cells without circulation.

In addition, we examined the kinetics of stem-like T cells to adopt a T_RM_ phenotype. To this end, after adoptive WT Pmel-1 transfer and B16F10 inoculation, we examined the mice either at an early time point when tumor size was less than 100mm^3^ or a late time point when tumor size reaches around 400mm^3^ (illustrated in **Supp. Fig. 4A**, tumor weight in **Supp. Fig. 4B**). As expected, significantly increased Pmel-1 T cells were detected in large tumor TDLN (**Supp. Fig. 4C**). Importantly, T_RM_ phenotype, especially the induction of CD69 on stem-like T cells was dramatically enhanced in late stage TDLN (**Supp. Fig. 4D** and **4E**). The percentage of CD69 on TDLN stem-like T cells was correlated very well with tumor size at early stage. After tumor weight reached around 300mg, T_RM_ phenotype on TDLN stem-like T cells was largely stabilized (**Supp. Fig. 4F**). This finding is highly consistent with the requirement of tumor antigen to induce T_RM_ phenotype on TDLN stemlike T cells.

To further validate this finding, we extended our analysis to bulk endogenous polyclonal CD8^+^ T cells. Indeed, we could consistently detect a small subset of Slamf6^+^Tcf-1^+^ endogenous CD8^+^ T cells, presumably representing stem-like CD8^+^ T cells (**Supp. Fig. 5A**). A substantial portion of these stem-like T cells, but not other endogenous CD8^+^ subsets, expressed both CD103 and CD69 (**Supp. Fig. 5A**). To be noted, Tcf-1^+^Slamf6^-^ subset likely included non-specific naïve CD8^+^ T cells that expressed CD103, but not CD69. Importantly, identical to donor Pmel-1 T cells (**Fig. 4D** and **4E**), when comparing stem-like endogenous CD8^+^ T cells isolated from different lymphoid organs, there was a stepwise reduction of T_RM_ phenotype from TDLN, NDLN to spleen (**Supp. Fig. 5B**).

Together, we have demonstrated that naturally primed tumor-specific CD8^+^ T cells differentiate into T_RM_s in TDLN in a TGF-β- and tumor antigen-dependent manner.

### Dynamic regulation of stem-like T cells in TDLN after tumor vaccine

Next, we focused on the response of TDLN stem-like T cells to tumor vaccine. To perform side-by-side comparison of WT vs *Tgfbr2^-/-^* T cells isolated from the same tissue of the same animal, we employed adoptive co-transfer system. As illustrated in **Fig. 5A**, naïve Pmel-1 T cells were isolated from congenically distinct WT and *Tgfbr2^-/-^* mice, mixed at a 1:1 ratio and adoptively co-transferred into C57BL/6 recipients followed by s.c. B16 tumor inoculation. Tumor vaccine was administrated similarly at a late stage. The vast majority of TDLN WT Pmel-1 T cells exhibited a stem-like phenotype before vaccination (**Fig. 5B** day 0). Lack of TGF-β signaling moderately reduced the portion of stem-like T cells at base line (**Fig. 5B** day 0). After tumor vaccine, stem-like T cells were slightly decreased for WT Pmel-1 T cells. In contrast, *Tgfbr2^-/-^* Pmel-1 T cells exhibited a much more dramatic reduction of stem-like subset after vaccination (**Fig. 5B**), consistent with enhanced differentiation of stem➔migratory effectors in *Tgfbr2^-/-^* cells.

**Figure 5.**
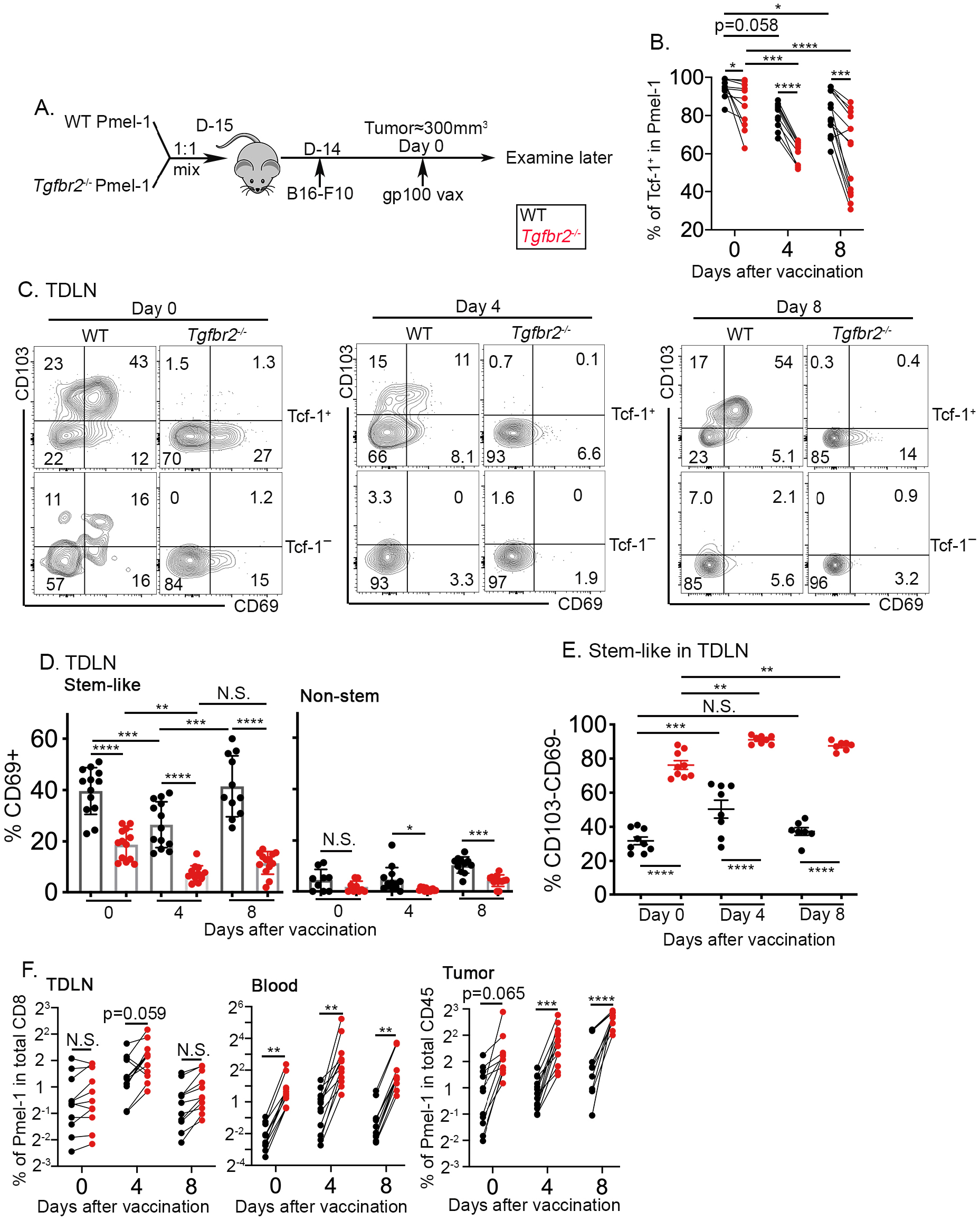
Tumor vaccine induces the loss of tissue residency in TDLN for stem-like T cells, accompanied by further differentiation into migratory effectors. (**A**) Experimental design. (**B**) The percentage of Tcf-1^+^ stem-like subset in TDLN Pmel-1 T cells at different timepoints post vaccination are shown. (**C**) Representative FACS profiles of pre-gated Tcf-1^+^ (Upper, stem-like) and Tcf-1^-^ (Lower, non-stem) Pmel-1 T cells in TDLN at different time points after vaccination are shown. (**D**) The percentage of CD69^+^ cells in stem-like (Left) and non-stem (Right) Pmel-1 T cells isolated from TDLN are shown. (**E**) The percentage of non-T_RM_ (CD69^-^CD103^-^) cells in stem-like Pmel-1 T cells isolated from TDLN is shown. (**F**) The percentage of donor Pmel-1 T cells in total CD8^+^ (**Left**, TDLN; **Middle**, blood) or total CD45^+^ cells (**Right**, tumor) are shown. Each pair of symbols in (B) and (F), each symbol in (D) and (E) represents the results from an individual recipient mouse. Pooled results from 4 (B) or 3 (D-F) independent experiments are shown. N.S., not significant, *, p<0.05, **, p<0.01, ***, p<0.001 and ****, p<0.0001 by Ordinary one-way ANOVA with multi-comparison posttest.

Importantly, this dynamic regulation of stem-like subset in response to tumor vaccine is TDLN-specific. In non-draining lymph nodes (NDLN), we did not detect any significant changes in the percentage of stem-like subset (**Supp. Fig. 6A**). In the spleen, we did find a reduction of stem-like subset in *Tgfbr2^-/-^* T cells compared with WT counterparts. However, we did not discover tumor vaccine-induced alterations (**Supp. Fig. 6B**).

Together, we find that in response to tumor vaccine, TDLN stem-like T cells differentiate into non-stem effectors and TGF-β inhibits this differentiation process.

### Tumor vaccine induces transient loss of T_RM_ phenotype in TDLN stem-like T cells

Then, we focused on the impacts of tumor vaccine on T_RM_ stem-like T cells in TDLN. Consistent with **Fig. 4**, we found that before vaccine administration, a significant portion of stem-like T cells exhibited a T_RM_ phenotype for WT Pmel-1 T cells while the expression of CD103 was completely abolished and that of CD69 was significantly reduced in *Tgfbr2^-/-^* ones (**Fig. 5C** left and **5D** left). Shortly after vaccination (i.e., day 4), WT stem-like Pmel-1 T cells carried greatly reduced T_RM_ markers while *Tgfbr2^-/-^* stem-like ones almost completely lost CD69 expression (**Fig. 5C** left vs middle and **5D** left). Day 8 after vaccination, WT stem-like T cells largely regained T_RM_ markers while *Tgfbr2^-/-^* ones still carried reduced level of CD69 (**Fig. 5C** left vs right and **5D** left). If we could define CD69^-^ CD103^-^ Tcf-1^+^ cells as non-T_RM_ stem-like cells, this subset was transiently induced in WT Pmel-1 T cells by tumor vaccine (**Fig. 5E**). In contrast, *Tgfbr2^-/-^* Pmel-1 T cells carried significantly increased population of non-T_RM_ stem-like T cells before tumor vaccine. Importantly, tumor vaccine further boosted sustained elevation of non-T_RM_ stem-like subset for *Tgfbr2^-/-^* cells (**Fig. 5E**). Tcf-1^-^ effectors largely exhibited a non-T_RM_ migratory phenotype in TDLN, especially after tumor vaccine for both WT and *Tgfbr2^-/-^* cells (**Fig. 5C** bottom row and **5D** right).

The greatly reduced T_RM_ phenotype in TDLN *Tgfbr2^-/-^* cells was translated into altered distribution. *Tgfbr2^-/-^* Pmel-1 T cells only exhibited a subtle increase in TDLN compared with co-transferred WT counterparts in the presence or absence of tumor vaccine (**Fig. 5F** left). In stark contrast, *Tgfbr2^-/-^* Pmel-1 T cells were the dominant population detected in the blood (At day 0, 12±4.4% WT vs 88±4.4% *Tgfbr2^-/-^*; day 4, 14±10% WT vs 86±10% *Tgfbr2^-/-^* day 8, 11±2.4% WT vs 89±2.4% *Tgfbr2^-/-^* and **Fig. 5F** middle). The dramatically increased circulation and migration of *Tgfbr2^-/-^* Pmel-1 T cells led to markedly increased accumulation inside tumor, especially after tumor vaccination (**Fig. 5F** right).

Interestingly, we could also detect similar tumor vaccine-induced changes of T_RM_ phenotype in Pmel-1 T cells isolated from non-draining LNs (NDLN) (**Supp. Fig. 7A**). However, we could not detect similar changes in either spleen (**Supp. Fig. 7B**) or tumor (**Supp. Fig. 7C**).

Together, tumor vaccine induces transient loss of T_RM_ phenotype in stem-like T cells in TDLN, which coincides with the differentiation from stem-like T cells into migratory effectors. *Tgfbr2^-/-^* CD8^+^ T cells carried dramatically reduced T_RM_ phenotype at base line in TDLN. *Tgfbr2^-/-^* stem-like CD8^+^ T cells exhibited enhanced and prolonged response to tumor vaccine, i.e., increased differentiation into migratory effectors.

### WT Pmel-1 T cells are highly enriched for T_RM_ gene signature

To further confirm the T_RM_ identity of stem-like T cells in TDLN, we FACS sorted WT and *Tgfbr2^-/-^* Pmel-1 T cells from TDLN and tumor after tumor vaccine. To eliminate the complication introduced by cell migration and focus on immediate local response induced by tumor vaccine, all samples were obtained from FTY720 treated animals. When WT and *Tgfbr2^-/-^* T cells from TDLN were compared, core T_RM_ signature(*58*) was significantly enriched in WT samples (**Fig. 6A**). In contrast, T_RM_ signature was not significantly enriched when WT tumor and *Tgfbr2^-/-^* tumor samples were compared (**Fig. 6C**). Specifically, the expression of a collection of T_RM_-associated genes, including *Itgae, Rgs10, Cdh1, Pmepa1, Skil* and *Ahr* was substantially reduced while the expression of circulating T cell signature genes *S1pr5* and *Klrg1* was enhanced in *Tgfbr2^-/-^* TDLN samples (**Fig. 6E**). Consistently, the expression of a panel of cell adhesion/cytoskeleton-related genes was dramatically differentiated between WT vs *Tgfbr2^-/-^* TDLN samples while showing similar patterns of expression in WT vs *Tgfbr2^-/-^* tumor samples (**Supp. Fig. 8C**). These results further validate that T_RM_/cell migration/ movement-associated genes represent the key difference between TDLN WT vs *Tgfbr2^-/-^* CD8^+^ T cells.

**Figure 6.**
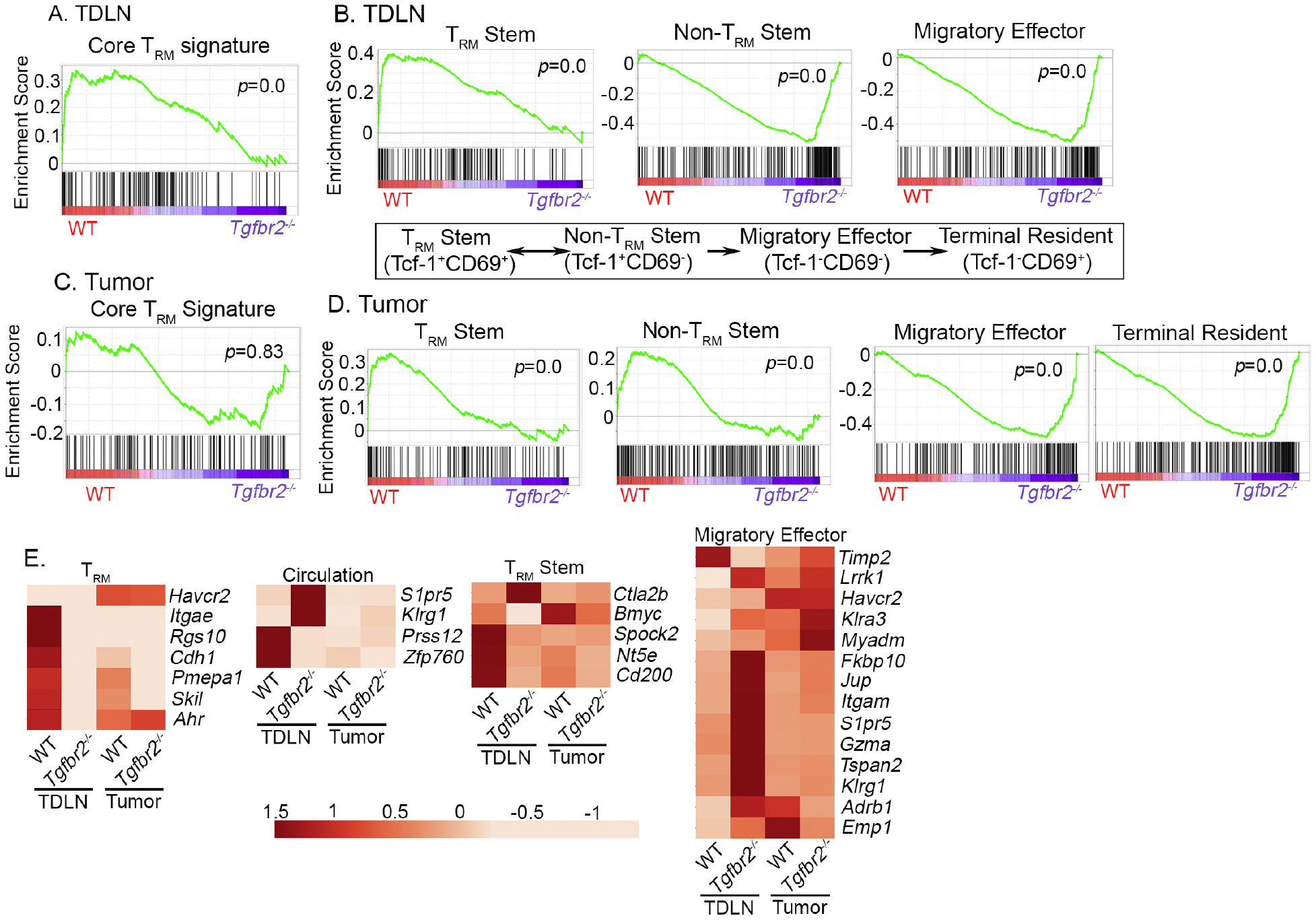
Enhanced differentiation from T_RM_ stem to non-T_RM_ stem in TDLN and accumulation of migratory effectors in tumor for *Tgfbr2^-/-^* Pmel-1 T cells. Bulk RNA-seq was performed on sorted Pmel-1 T cells isolated from TDLN and tumor. GSEA (gene set enrichment analysis) results are shown. (**A**) Core T_RM_ signature enrichment in TDLN samples. (**B**) The signatures of exhuasted CD8^+^ subsets in TDLN samples. Published differentiation pathway is illustrated in bottom panel. (**C**) Core T_RM_ signature enrichment in tumor samples. (**D**) The signatures of exhuasted CD8^+^ subsets in tumor samples. (**E**) Heatmap of differentially expressed signature genes. All groups contain biological independent duplicates.

Taking advantage of the recently established gene signatures of exhausted CD8^+^ T cell subsets from a mouse chronic viral infection model (*36*), we found that CD69^+^ Tcf-1^+^ (T_RM_ stem) signature was highly enriched in WT over *Tgfbr2^-/-^* samples from TDLN. In contrast, *Tgfbr2^-/-^* TDLN samples were highly enriched for CD69^-^Tcf-1^+^ (non-T_RM_ stem) and CD69^-^ Tcf-1^-^ (migratory effector) signatures (**Fig. 6B, 6E** and **Supp. Fig. 8C**). Interestingly, this pattern of gene signature enrichment was TDLN-specific, in tumor-derived Pmel-1 T cells, stem T cell signatures (both T_RM_ stem and non-T_RM_ stem) were enriched in WT while effector and terminally exhausted T cell signatures were enriched in *Tgfbr2^-/-^* samples (**Fig. 6D**), presumably due to the facts that T cell migration from TDLN to tumor was inhibited by FTY720 and enhanced differentiation from stem➔effector occurred inside tumor in the absence of TGF-β signaling. These results further support our conclusion that TDLN functions as a powerhouse for tumor vaccine and continuous migration from TDLN to tumor is required to sustain *Tgfbr2^-/-^* T cell response to tumor vaccine. To be noted, when comparing TDLN vs tumor samples, core T_RM_ gene signature was highly enriched in tumor samples for both WT and *Tgfbr2^-/-^* (**Supp. Fig. 8**), consistent with previous findings(*58*) and presumably due to the facts that terminally exhausted T cells (i.e., Tcf-1^-^ CD69^+^) carry highly enriched T_RM_ signature(*36*). Together, transcriptional profiling is largely consistent with our phenotypic analysis that tumor-specific CD8^+^ T cells differentiate into T_RM_ in TDLN in a TGF-β-dependent manner. In contrast, in tumor infiltrating T cells, T_RM_ signature may be controlled by both TGF-β-dependent and - independent pathways.

## Discussion

Here, we have demonstrated that tumor draining lymph nodes function as a reservoir for tumor-specific stem-like CD8^+^ T cells to reside. A large portion of stem-like T cells differentiate into T_RM_ in a tumor antigen- and TGF-β-dependent manner in TDLN. Importantly, loss of T_RM_ identity is required for the migration from TDLN to tumor and efficient response to tumor vaccine. For *Tgfbr2^-/-^* stem-like T cells, defective T_RM_ differentiation in TDLN leads to enhanced stem➔effector differentiation, elevated and sustained response to tumor vaccine. Importantly, our findings emphasize the importance of TDLN in tumor immunotherapies, i.e., TDLNs function as a reservoir to host T_RM_ stemlike T cells.

A large body of evidence has demonstrated that T_RM_ phenotype of tumor infiltrating T cells is often associated with improved tumor control and better outcomes(*51*). Considering the facts that TGF-β usually promotes T_RM_ differentiation and maintenance, it is challenging to completely explain why TGF-β inhibitors/blockers can synergize with tumor immunotherapies to improve anti-tumor immunity. Our results provide another perspective that the positive correlation of T_RM_ with tumor control may be a tumor-specific observation. TDLN-hosted T_RM_ is negatively associated with direct tumor killing. TDLN-targeted TGF-β blocking strategy may have the potential to be developed into a “universal adjuvant” for tumor vaccine. Further, our results imply that T_RM_ program may have a tissue/organ-specific component in terms of TGF-β dependency, which also favors the use of TGF-β inhibitors/blockers in tumor immunotherapies.

In contrast to previous findings that tumor vaccine promotes tumor control in WT mice, we did not find detectable delay of tumor progression in vaccinated WT Pmel-1 recipient mice. We believe the difference is due to the time when tumor vaccine is administrated. In previous research, peptide vaccine is often given when tumor is palpable. In contrast, we administrated tumor vaccine when tumor size reached around 300mm^3^. Indeed, when comparing early-vs late-stage TDLN, we found that T_RM_ marker CD69 expression was tightly associated with tumor size. This finding suggests that T_RM_ differentiation is a tumor-stage dependent feature and also provides an explanation why early vaccine can boost anti-tumor immunity in WT cells when tissue residency is not fully established in TDLN.

CD103 is another prominent TGF-β-dependent T_RM_ marker, especially for mucosal T_RM_s. In our system, even though the expression of CD103 was largely consistent with a T_RM_ marker, alternative explanation could not be completely ruled out. CD103 expression was not always associated with CD69 expression. For example, CD103^+^CD69^-^ stem-like T cells could be easily identified in the spleen or early stage TDLN. It has been suggested that CD103^+^CD8^+^ T cells represent a tolerant T cell subset and express Foxp3 at RNA level in a similar B16 tumor model(*59*). We could not detect any Foxp3 expression in Pmel-1 T cells at protein level in our system (not depicted). Alternatively, CD103^+^CD69^-^ cells may represent an intermediate stage of T_RM_ differentiation, similar to the observation of small intestine intra-epithelia lymphocyte (IEL) T_RM_ differentiation at an early stage after viral infection(*60*). The difference between CD103^+^CD69^-^ vs CD103^+^CD69^+^ stem-like T cells warrants future investigation. Together, we have established that stem-like CD8^+^ T cells differentiate into TDLN-resident T cells in a TGF-β and tumor antigen-dependent manner. Loss of TDLN residency is required for efficient response to tumor vaccine and differentiate into migratory effectors, which may represent another highly regulated step to be targeted for tumor immunotherapy.

## Materials and Methods

### Mice

C57BL/6J (B6) WT and Pmel-1 TCR transgenic mice (B6.Cg-Thy1^a^/Cy Tg (TcraTcrb) 8Rest/J) were obtained from the Jackson Laboratory. *Tgfbr2*^f/f^ dLck-cre OT-1 mice were described before (*13, 61*). *Tgfbr2*^f/f^ mice were originally from S. Karlsson (*62*) and dLck-cre mice were originally from N. Killeen (*63*). OT-1 mice were originally from Dr. Michael J. Bevan (University of Washington). All mice were housed at our specific pathogen-free animal facilities at the University of Texas Health at San Antonio (San Antonio, TX). All experimental mice have been backcrossed to C57BL/6 background for more than 12 generations. All experiments were done in accordance with the University of Texas Health Science Center at San Antonio Institutional Animal Care and Use Committee guidelines.

### Tumor Cell Lines

C57BL/6 derived melanoma lines B16F10 and B16-OVA were generous gifts from Dr. Tyler Curiel (UT Health San Antonio). Both lines were maintained in DMEM complete media (10% FBS + 1% L-glutamine, + 1% pen/strep + 0.1mM non-essential amino acids) at 37 °C in 5% CO_2_. All cell culture medium and supplements were purchased from Invitrogen.

### Naïve T Cell Isolation and Adoptive Transfer

Naïve CD8^+^ T cells (OT-1 or Pmel-1) were isolated from pooled spleen and lymph nodes using MojoSort™ mouse CD8 T cell isolation kit (Biolegend) following manufacturer’s instruction. During the first step of biotin antibody cocktail incubation, biotin-αCD44 (IM7, Biolegend) was added to label and deplete effector and memory T cells. Isolated naïve CD8^+^ T cells were numerated, 1:1 mixed when indicated, 10^5^ cells adoptively transferred into each sex-matched unmanipulated B6 recipient via an i.v. route.

### Tumor Inoculation and Immunotherapies

One day after naïve CD8^+^ T cell transfer, the flank of mice was shaved and B16F10 or B16-OVA cells (2.5~3×10^5^) were mixed with Matrigel (Corning, final concentration 5mg/ml) and injected subcutaneously (s.c.). Tumor volumes were estimated by measuring the tumor size in three dimensions (length, width and height) using a caliper. The tumor volume was calculated according to the formula (π/6 x length x width x height). Mice were sacrificed at the indicated time points or when the estimated tumor volume reached > 1000mm^3^ (endpoint) and the weight of the excised tumor mass was determined.

For tumor vaccination, mice were injected subcutaneously (beside the tumor) with 50 μg/mouse poly(I:C) (Invivogen) together with antigenic peptides (for Pmel-1, gp100_25-33_; for OT-1, OVA_257-264_) at 10μg/mouse (peptides were purchased from Anaspec). The control (CON) group mice were injected with PBS in same volume.

For checkpoint blockade, mice were injected with rat anti-mouse PD-L1 or isotype control (BioXcell) (200 μg per injection, i.p.) once every four days for a total of three injections.

### FTY720 Treatment

FTY720 (ENZO Life Sciences), stock solution (4mg/mL in DMSO) was diluted to 100 μg/mL in double distilled water (dd water) directly before administration and 25 μg/mouse were applied daily by oral gavage for the duration of the experiment. Water containing same concentration of DMSO was used as control.

### Lymphocyte Isolation

Blood, spleen, TDLN (inguinal LN on tumor side), NDLN (inguinal LN opposite tumor side, superficial cervical LN, axillary LN) and tumor were isolated. CD8^+^ T cells from spleen and LN were obtained by mashing through a 100 μm nylon cell strainer (BD Falcon). Red blood cells were lysed with a hypotonic Ammonium-Chloride-Potassium (ACK) buffer (prepared in the lab). For TIL isolation, tumors were excised and digested with 1 mg/mL collagenase B (Roche) and 0.02 mg/mL DNaseI (D5025 from Sigma) at 37°C for 45min. Digested tumors were mashed through 70 μm filters. Single cell suspensions were then resuspended in RPMI1640 complete media (10% FBS + 1% L-glutamine, + 1% pen/strep + 0.1mM non-essential amino acids) for flow cytometry.

### Antibodies and Flow cytometry

Fluorescence dye labeled antibodies specific for PD-1 (J43), CD8β (H35-17.2), Granzyme A (GzA-3G8.5), Granzyme B (GB11), CD45.1 (A20), CD45.2 (104), CD8α (536.7), CD69 (H1.2F3), CD103 (2E7), CD39 (24DMS1), CD101 (Moushi101), CX3CR1 (SA011F11), Slamf6 (330-AJ) were purchased from eBioscience, Biolegend, Invitrogen and Tonbo. Anti-CD16/32 (2.4G2) was produced in the lab and used in all FACS staining as FcR blocker. Intranuclear staining for Tcf-1 was performed using a Foxp3 staining buffer set (Tonbo bioscience) and stained with anti-Tcf-1 (C63D9, Cell Signaling). Intracellular staining for Granzyme A and Granzyme B was performed using permeabilization buffer (Invitrogen) following fixation. For intracellular cytokine staining, freshly isolated lymphocytes from tumor were cultured in complete RPMI in the presence of Golgi STOP (BD) with αCD3 (1μg/ml, 2C11, BioXcell) +αCD28(1μg/ml, E18, Biolegend) for 4 hours. Stimulated cells were surface stained, fixed, permeablized and intracellular stained by anti-IFN-γ antibody (XMG1.2, Biolegend) and anti-TNF antibody (MP6-XT22, Biolegend). Ghost Dye™ Violet 510 (Tonbo bioscience) was used to identify live cells. For some tumor samples, fluorescent counting beads (AccuCount Fluorescent Particles from Spherotech) were added before analysis to calculate the number of donor Pmel-1 CD8^+^ T cells. Washed and fixed samples were analyzed by BD LSRII or BD FACSCelesta, and analyzed by FlowJO (TreeStar) software.

### Gene Expression Profiling

Day 4 after tumor vaccine, TDLNs and tumors containing WT and *Tgfbr2^-/-^* Pmel-1 T cells were dissected. To limit T cell migration, FTY720 was administrated. Donor Pmel-1 T cells were FACS sorted based on CD8, CD45.1 and CD45.2. RNA was extracted from sorted cells using the Quick-RNA™ MiniPrep, according to manufacturer’s instructions (Zymo Research). RNA-seq analysis was performed by Novogene. Original RNA-seq results can be accessed by GSE176525.

### Statistical Analysis

Ordinary One-way ANOVA, Mantel-Cox or Student *t*-test from Prism 9 was used.

## Acknowledgements

We thank Dr. Tyler Curiel for providing B16F10 and B16-OVA cell lines. We thank Dr. Ananda Goldrath for discussion and suggestions. This work is supported by NIH grants AI125701 and AI139721, Cancer Research Institute CLIP program and American Cancer Society grant RSG-18-222-01-LIB to N.Z. We thank Karla Gorena and Sebastian Montagnino for FACS sorting. Data generated in the Flow Cytometry Shared Resource Facility were supported by the University of Texas Health Science Center at San Antonio (UTHSCSA), NIH/NCI grant P30 CA054174-20 (Clinical and Translational Research Center [CTRC] at UTHSCSA), and UL1 TR001120 (Clinical and Translational Science Award).

---

**Supplemental Figure 1.**
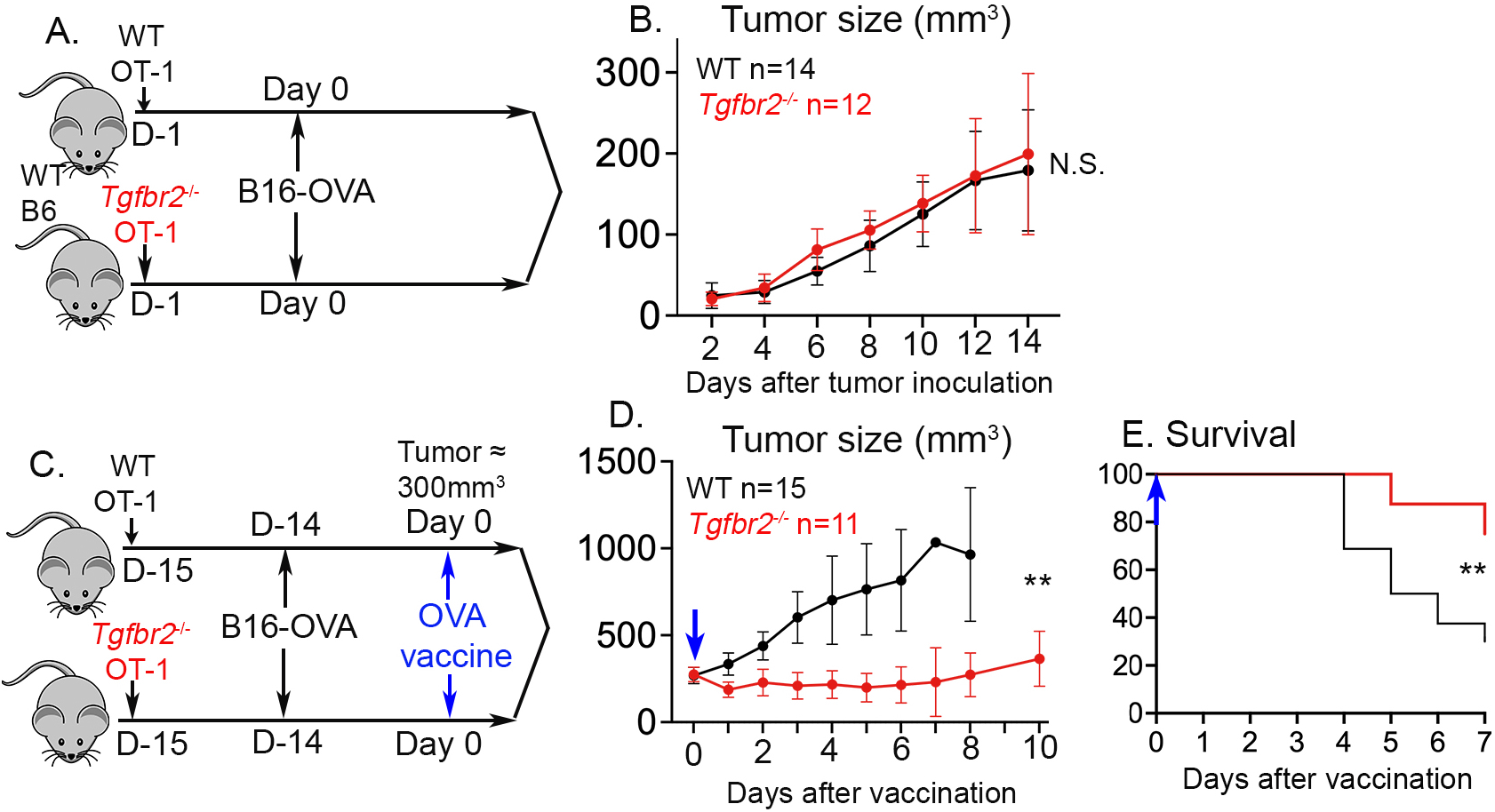
*Tgfbr2^-/-^* OT-1 T cells synergize with OVA peptide vaccination to control B16-OVA. (**A**) and (**B**), OT-1 transfer alone. (**A**) Schematics; (**B**) Tumor growth. (**C**) to (**E**), OT-1+OVA vaccine. (**C**) Schematics; (**D**) Tumor growth and (**E**) Survival curve after vaccination are shown. Pooled results from 2-3 independent experiments are shown. N.S., not significant, **, p<0.01 by Student *t*-test or Mantel-Cox test.

**Supplemental Figure 2.**
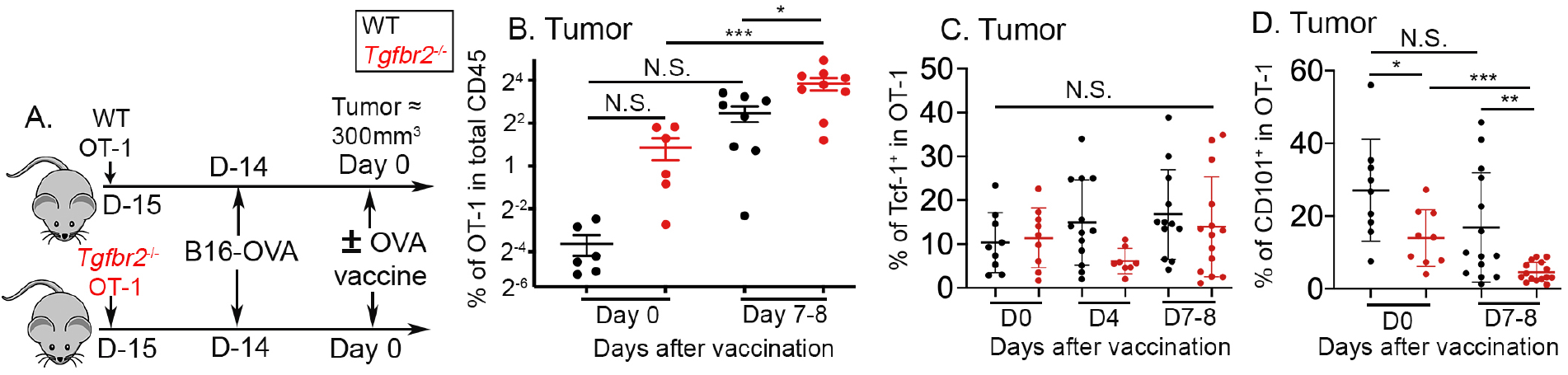
Enhanced accumulation of *Tgfb2^-/-^* OT-1 T cells inside tumor after vaccination. (**A**) Schematics. (**B**) The percentage of OT-1 T cells in total CD45^+^ tumor infiltrating cells is shown. (**C**) The percentage of Tcf-1^+^ subset in tumor infiltrating OT-1 T cells is shown. (**D**) The percentage of CD101^+^ subset in tumor infiltrating OT-1 T cells is shown. Each symbol represents the results from an individual mouse. Pooled results from 3 independent experiments are shown. N.S., not significant, *, p<0.05, **, p<0.01 and ***, p<0.001 by Ordinary one-way ANOVA with multi-comparison posttest.

**Supplemental Figure 3.**
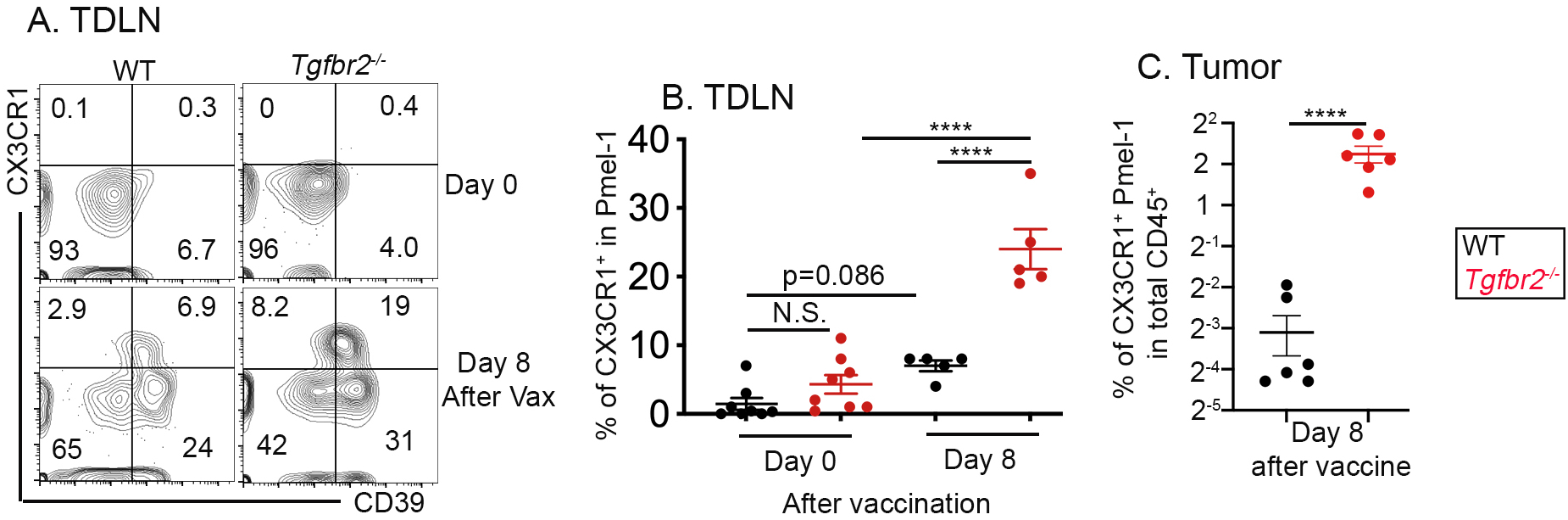
Tumor vaccine boosts the differentiation of CX3CR1^+^ migratory effectors in TDLN and tumor. Similar experimental setup as in Fig. 2A. (**A**) Representative FACS profiles of donor Pmel-1 T cells in TDLN before and after tumor vaccine are shown. (**B**) The percentage of CX3CR1^+^ effectors in Pmel-1 T cells from TDLN is shown. (**C**) The percentage of CX3CR1^+^ Pmel-1 T cells in total CD45^+^ tumor infiltrating cells after tumor vaccine is shown. Pooled results from 2 independent experiments are shown. Each symbol in (B) and (C) represents the results from an individual mouse. N.S., not significant and ****, p<0.0001 by Ordinary one-way ANOVA with multi-comparison posttest (B) or Student *t*-test (C).

**Supplemental Figure 4.**
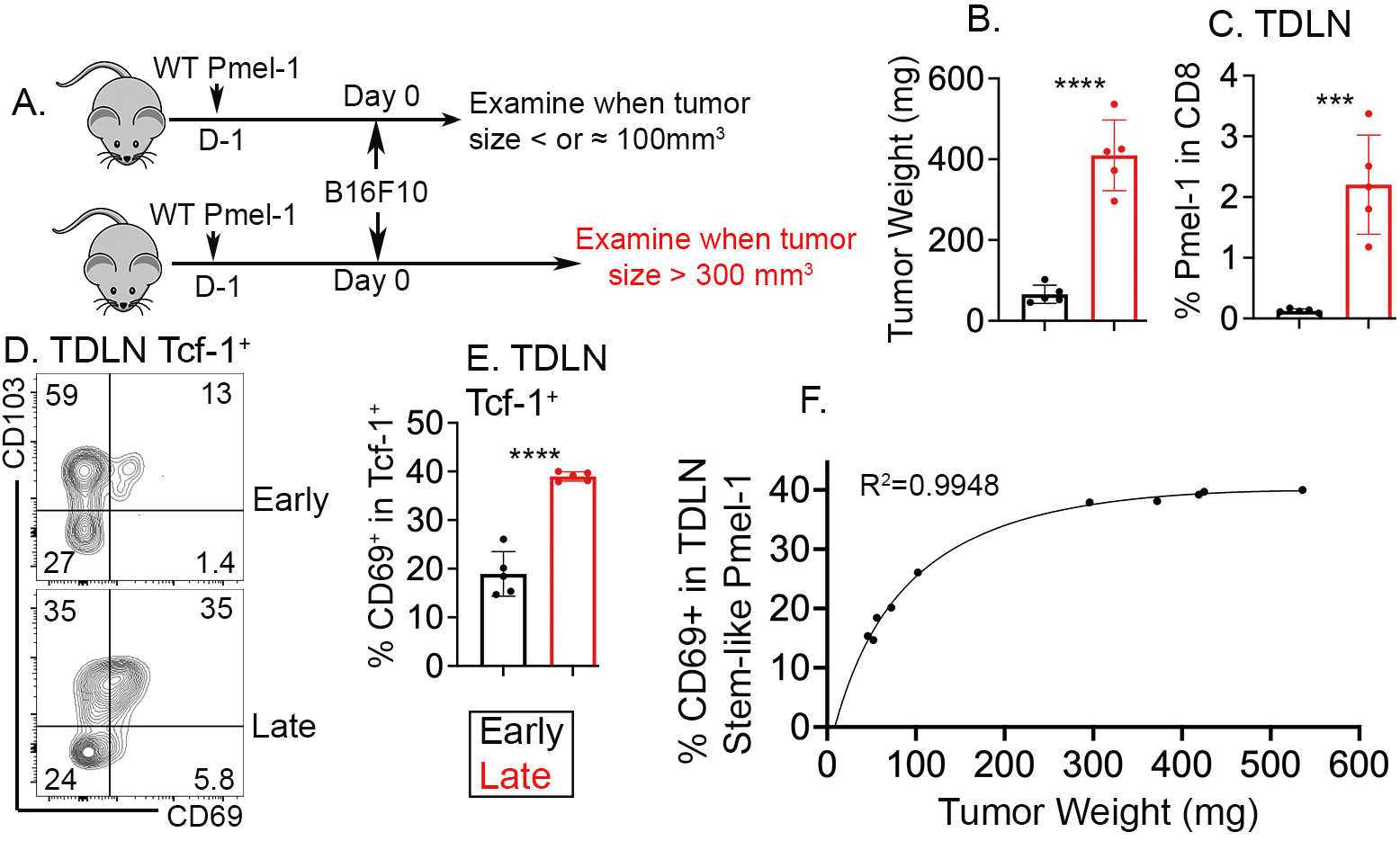
Tumor size-dependent accumulation of T_RM_ stem-like T cells in TDLN. (**A**) Experimental design. Briefly, WT Pmel-1 T cells were adoptively transferred into B6 recipients followed by B16F10 s.c. inoculation. At 9-10 days later (**early**) or 19-20 days later (**late**), tumor size and TDLN were examined. (**B**) Tumor weight for early vs late time points is shown. (**C**) The percentage of Pmel-1 T cells in total CD8 in TDLN is shown. (**D**) Representative FACS profiles of pre-gated Tcf-1^+^ Pmel-1 T cells from TDLN are shown. (**E**) The percentage of CD69^+^ subset in Tcf-1^+^ Pmel-1 T cells isolated from TDLN is shown. (**F**) Nonlinear regression of the percentage of CD69^+^ among Tcf-1^+^ Pmel-1 T cells in TDLN vs tumor weight of the same animal. Each symbol in (B), (C), (E) and (F) represents the results from an individual mouse. ***, p<0.001 and ****, p<0.0001 by Student *t*-test.

**Supplemental Figure 5.**
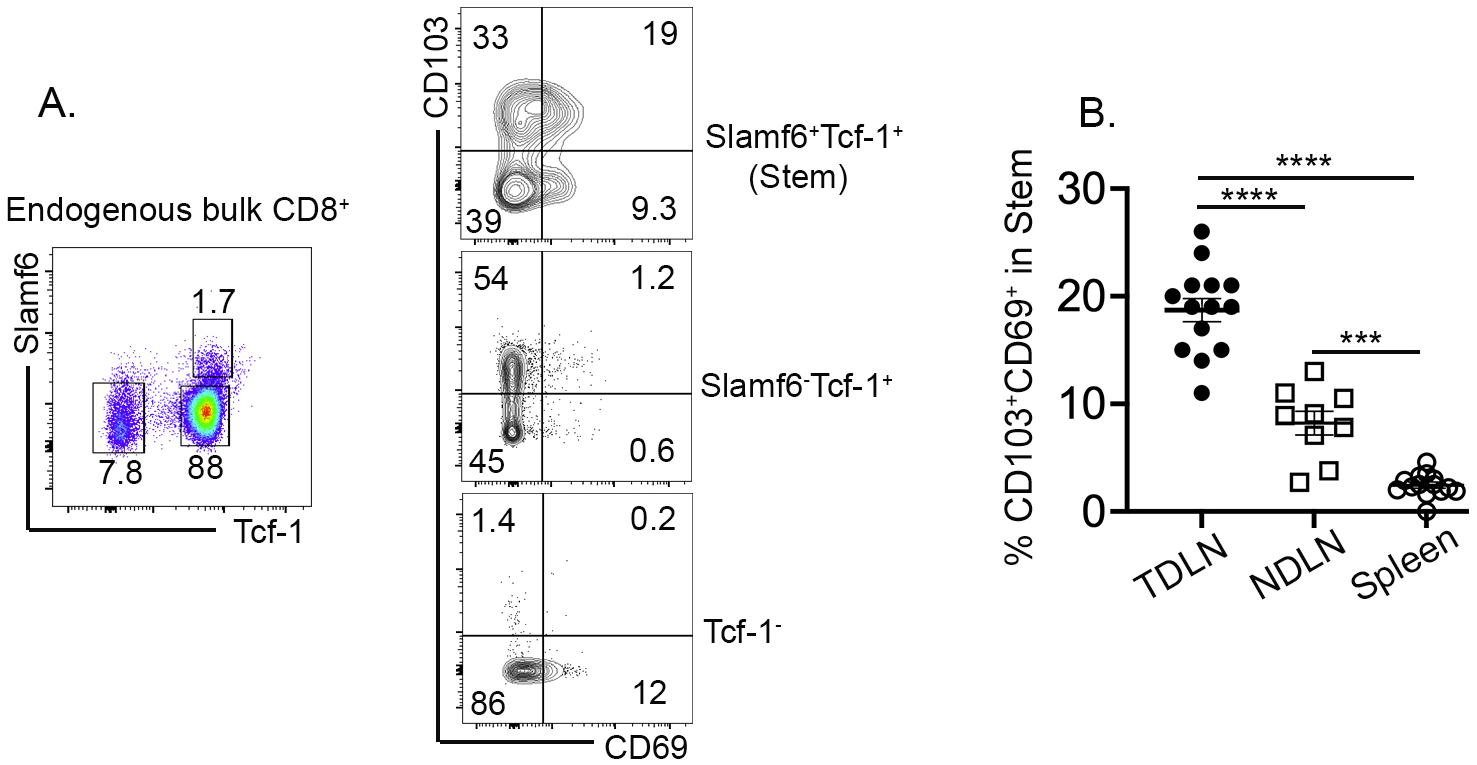
Stem-like endogenous polyclonal CD8^+^ T cells differentiate into T_RM_ in TDLN. Same experimental setup as in Fig. 4A. (**A**) Left, representative gating strategy on endogenous bulk CD8^+^ T cells isolated from TDLN; Right, representative FACS profiles of pre-gated endogenous CD8^+^ T cells to show T_RM_ phenotype. (**B**) The percentage of CD69^+^CD103^+^ subset in endogenous stem-like CD8^+^ T cells isolated from different lymphoid organs. Each symbol in (B) represents the results from an individual mouse. Pooled results from 3 independent experiments are shown. ***, p<0.001 and ****, p<0.0001 Ordinary one-way ANOVA with multi-comparison posttest.

**Supplemental Figure 6.**
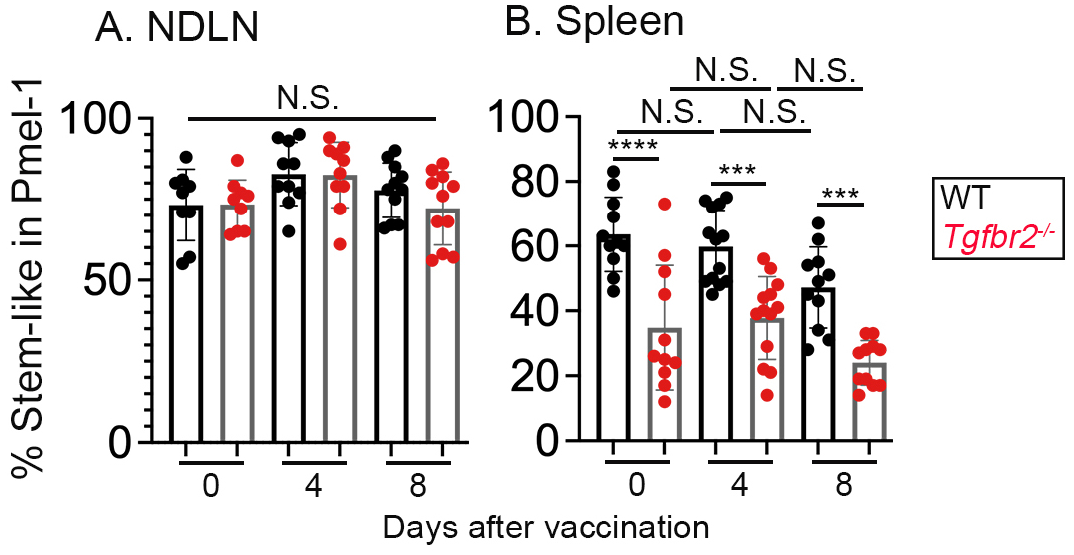
Stem-like T cell subset in other lymphoid organs after tumor vaccination. Same experimental setup as in Fig. 5. The percentage of stem-like subset in donor Pmel-1 T cells isolated from NDLN (**A**) and spleen (**B**) at different time points after vaccination are shown. Pooled results from 3 independent experiments are shown. Each symbol represents the results from an individual recipient mouse. N.S., not significant, ***, p<0.001 and ****, p<0.0001 by Ordinary one-way ANOVA with multicomparison posttest.

**Supplemental Figure 7.**
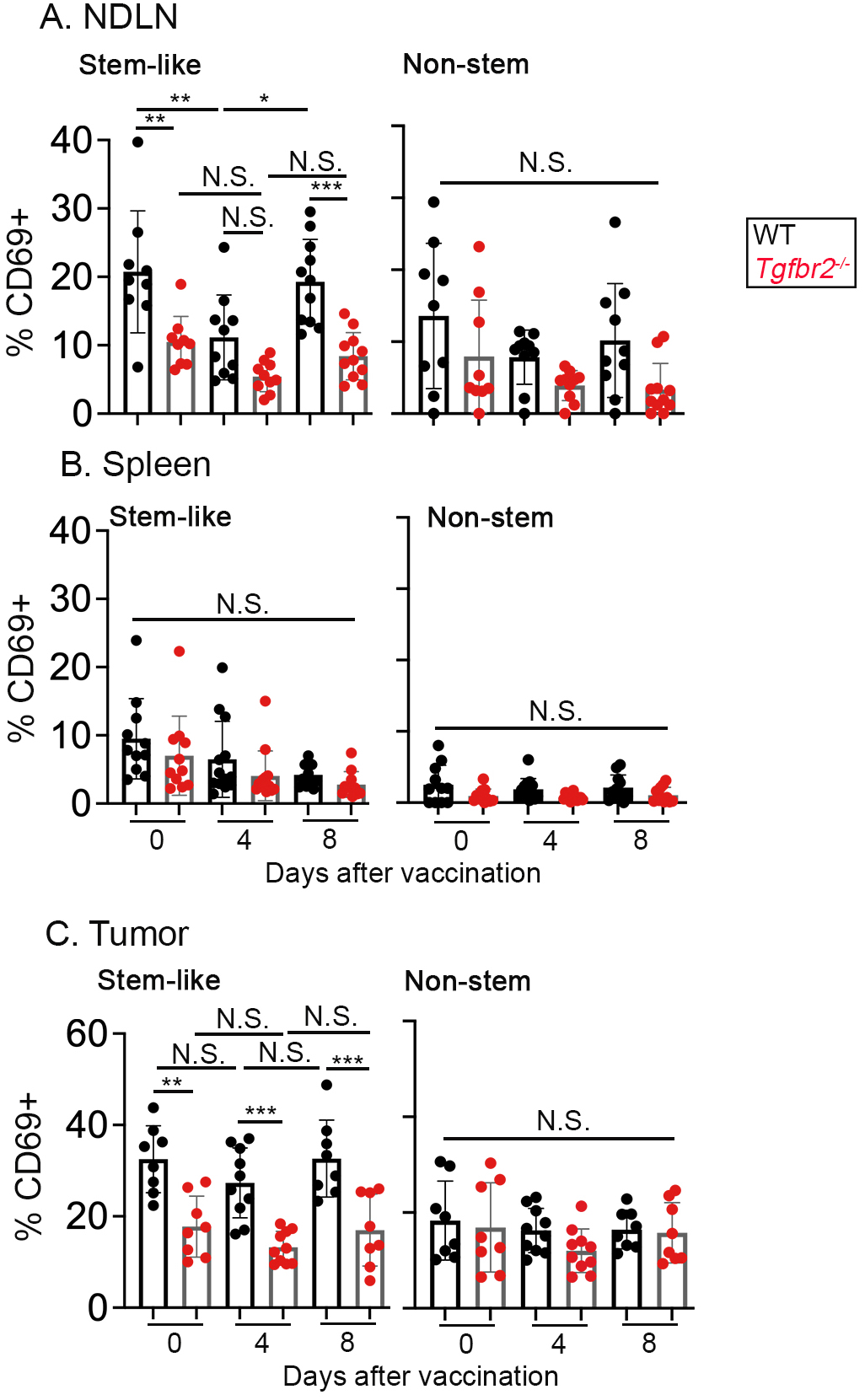
The alteration of tissue residency after tumor vaccination. Same experimental setup as in Fig. 5. (**A**) The percentage of CD69^+^ cells in stem-like (Left) and non-stem (Right) Pmel-1 T cells isolated from NDLN are shown. (**B**) The percentage of CD69^+^ cells in stem-like (Left) and non-stem (Right) Pmel-1 T cells isolated from spleen are shown. (**C**) The percentage of CD69^+^ cells in stem-like (Left) and nonstem (Right) Pmel-1 T cells isolated from tumors at different time points after vaccination are shown. Pooled results from 3 independent experiments are shown. Each symbol represents the results from an individual recipient mouse. N.S., not significant, *, p<0.05, **, p<0.01 and ***, p<0.001 by Ordinary one-way ANOVA with multi-comparison posttest.

**Supplemental Figure 8.**
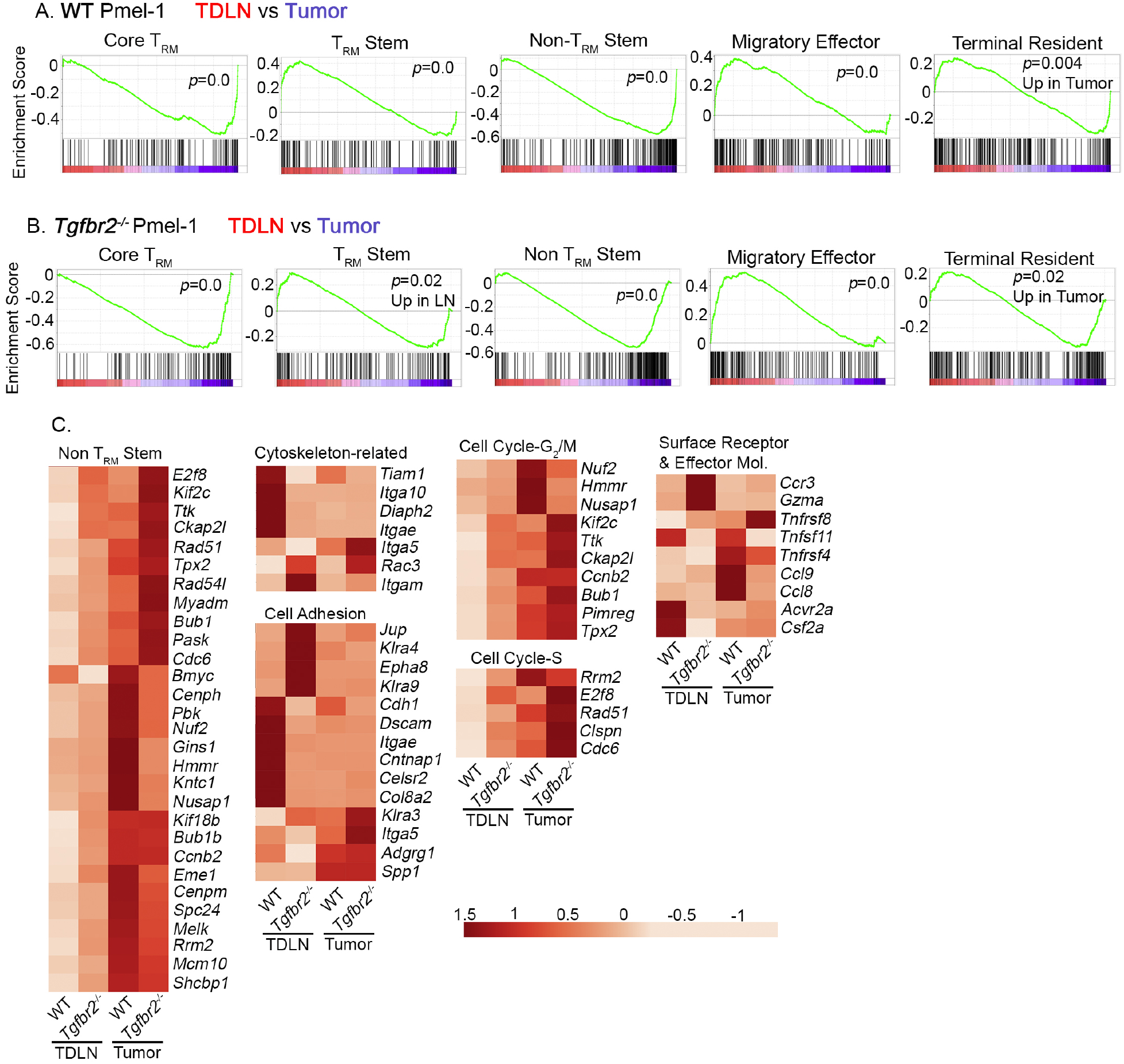
Enrichment of T_RM_ signature genes in tumor. GSEA results comparing TDLN vs tumor samples. (**A**) WT and (**B**) *Tgfbr2^-/-^* samples (**C**) Heatmap of DEGs for selected signature genes.

